# NREM sleep brain networks modulate cognitive recovery from sleep deprivation

**DOI:** 10.1101/2024.06.28.601285

**Authors:** Kangjoo Lee, Yimeng Wang, Nathan E. Cross, Aude Jegou, Fatemeh Razavipour, Florence B. Pomares, Aurore A. Perrault, Alex Nguyen, Ümit Aydin, Makoto Uji, Chifaou Abdallah, Birgit Frauscher, Habib Benali, Thien Thanh Dang-vu, Christophe Grova

## Abstract

Decrease in cognitive performance after sleep deprivation and its recovery following sleep suggests the key role of sleep, especially non-rapid eye movement (NREM) sleep, in maintaining cognition. It remains unknown whether brain network reorganization in NREM sleep stages N2 and N3 can uniquely be mapped onto individual differences in cognitive performance after a recovery nap following sleep deprivation. Using resting state functional magnetic resonance imaging (fMRI), we quantified the integration and segregation of brain networks during N2 and N3 sleep in a 1-hour nap following 24-hour sleep deprivation, compared to well-rested wakefulness. Here, we introduce a new analytic framework called the hierarchical segregation index (HSI) to quantify network segregation across spatial scales, from whole-brain to the voxel level, by identifying spatio-temporally overlapping large-scale networks and the corresponding voxel-to-region hierarchy. We found that network segregation increased in the posterior cingulate cortex during both N2 and N3 sleep compared to wakefulness. In contrast, increased segregation in the visual cortex and decreased segregation in the posterior ventromedial prefrontal cortex during N2 returned to levels comparable to wakefulness during N3. More segregation during N3 sleep was associated with worse recovery of working memory, executive attention, and psychomotor vigilance after the nap. Sleep-related changes in network segregation emerged at different spatial scales, with regional- and voxel-level alterations being associated with different cognitive tasks. Finally, we demonstrated the sensitivity and reliability of voxel-level HSI to provide key insights into within-region variation, suggesting a mechanistic understanding of how NREM sleep replenishes cognition after sleep deprivation.

## Introduction

Sleep stages, including non-rapid eye-movement (NREM) sleep stages, are defined following a polysomnography exam, using notably spontaneous brain oscillations detected by the scalp electroencephalography (EEG) (1). NREM sleep stage 2 (N2) is characterized by transient oscillations called spindles that play a role in memory consolidation and in the protection of sleep quality (2, 3). NREM sleep stage 3 (N3) is characterized by slow waves, being related to sleep homeostasis (4). Using simultaneous recordings of scalp EEG with functional magnetic resonance imaging (fMRI), one can monitor hemodynamic processes through the measurement of the blood-oxygen-level-dependent (BOLD) signals, during different sleep stages (5–7).

Compared to resting state networks that are consistently found during wakefulness (8), EEG-fMRI studies reported that functional connectivity decreases during sleep within the default mode network (9), between the default mode and its anti-correlated networks (10, 11), within the frontoparietal network (12, 13) and between the thalamus and cortical areas (14). Using high-density scalp EEG, neuronal activation of the premotor region evoked by transcranial magnetic stimulation (TMS) propagated to neighboring regions in the awake state (15). During NREM sleep, this evoked response disappeared more rapidly with no propagation, suggesting less involvement of long distance connections while prioritizing local segregated activity (15). Evidence converges to suggest sleep-dependent brain-wide changes in functional integration, however, specific network reorganization during N2 and N3 and their roles in the maintenance of normal cognitive functions remain unclear.

Cognitive processes involve neural circuits recruiting distributed regions across the cerebral cortex. Sleep plays an important role in the maintenance and recovery of cognitive performances. Reduced consciousness during NREM sleep was associated to increased segregation within six large-scale cortical networks during NREM sleep when compared to wakefulness (16). To investigate patterns of segregation within nested hierarchical brain networks, Boly et al. (16) applied a metric entitled the functional clustering ratio (FCR), derived from information theory measures (17). FCR calculates the ratio between within- and between-network integration, quantifying to what extent a large-scale network segregates into non-overlapping smaller modules. Recently, in Cross et al., we estimated FCR from task fMRI data acquired in a three-day protocol to experimentally manipulate vigilance states (**Fig. 1A**), including 24 hours of sleep deprivation (SD) and one-hour recovery sleep after SD (12). In this study, we compared FCR estimated from task fMRI during well-rested (WR) awake state, task fMRI during sleep-deprived (SD) state, resting state fMRI during NREM sleep after SD, and task fMRI data during a post-recovery nap (PRN) condition. After regressing out the effect of the task, we found increased FCR (segregation) during NREM sleep when compared to WR and SD states, which then decreased after the nap (WR*<*PRN*<*SD*<*NREM) (12). Moreover, individuals exhibiting more segregation (increased FCR) in task fMRI during SD state compared to resting baseline showed worse task performance with SD, while individuals exhibiting more integration in task fMRI during PRN when compared to SD (PRN*<*SD) showed better recovery of task performances (12). However, specific patterns of network segregation during NREM sleep in relation to cognition remain unexplored, especially when considering specifically N2 and N3 sleep stages separately.

**Fig. 1.**
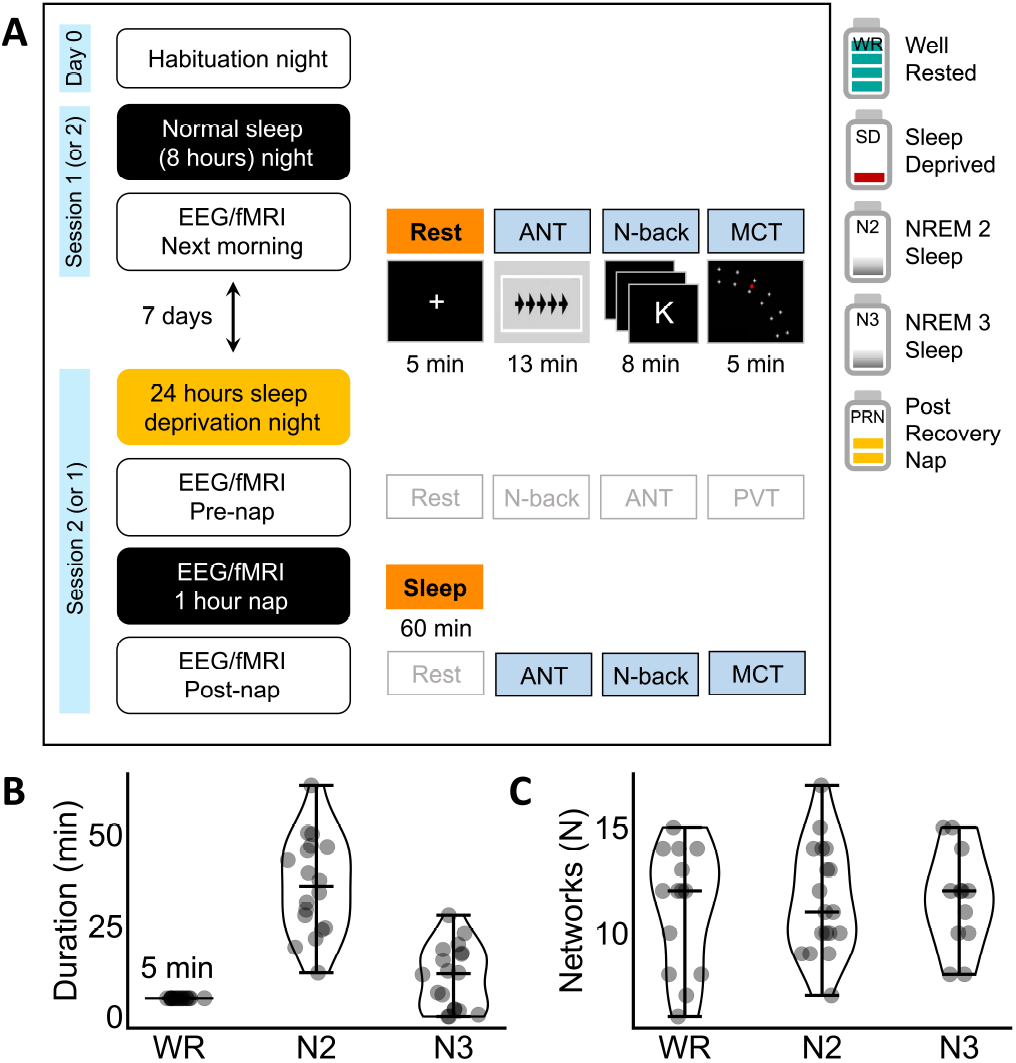
Simultaneous high-density EEG/fMRI data to assess the impact of sleep deprivation. **(A)** Overview of study design (12). Participants visited the lab three times: a habituation night to evaluate their eligibility, one night with normal sleep and one night with total SD followed by a one-hour recovery nap inside the MRI scanner. Cognitive tasks (ANT: Attention network task (22) for testing executive attention, N-back (23) for working memory, and MCT: Mackworth clock test (24) for psychomotor vigilance) were applied three times: after the normal night of sleep, before and after the recovery nap following SD. Data analyzed in this study are highlighted in orange (imaging data) and blue (behavioral data) boxes. Task fMRI were not used in this study (see (12)). Resting state fMRI at pre-nap and post-nap were not used because many subjects fell asleep during the scan. EEG was used only for sleep stage identification. **(B)** Total duration of WR (5 min) as well as N2 and N3 sleep during the one-hour nap after SD of each individual considered for the analysis. **(C)** Preserved distribution of the global network scale, i.e., the total number (*N*) of networks, across states. *N* was estimated for each 5-minute individual fMRI run using SPARK (19, 20).

To fill this gap, we investigate network segregation of resting state fMRI during N2 and N3 sleep stages separately in comparison to WR using the same dataset as in (12) (**Fig. 1**), in relation to cognitive recovery after the sleep. We hypothesize that network reorganization during NREM sleep is associated with the level of recovery of cognitive performances after the recovery nap following SD. Studying patterns of segregation during different NREM stages at finer spatial resolutions than at the regional or network level may help to better understand how NREM sleep replenishes cognition after SD. Here, to quantify functional network segregation at the voxel level, we propose a new metric, the hierarchical segregation index (HSI). The HSI is defined using a novel data-driven voxel-based measure of functional integration, *k*-hubness, assessing the number of overlapping functional networks involved in a voxel (**Fig. 2**). *k*-hubness is estimated using a recently developed method for sparsity-based analysis of reliable *k*-hubness (SPARK) based on sparse dictionary learning of individual fMRI (18–21). We identify the patterns of network segregation characterizing the overall NREM sleep and separate unique patterns of functional networks specific to N2 and N3 sleep, in relation to the recovery of cognitive task performances after the recovery nap following SD.

**Fig. 2.**
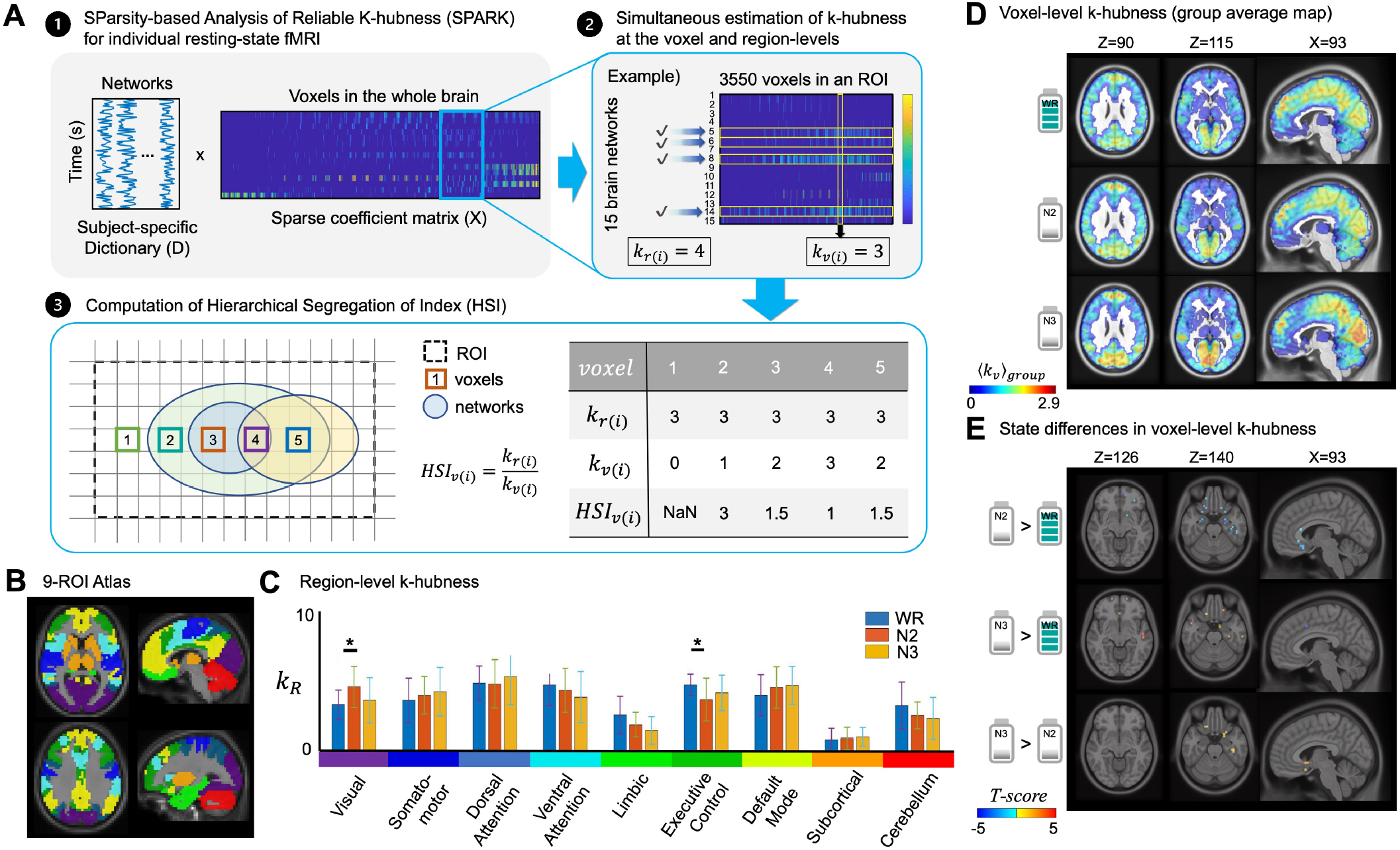
Definition and estimation of the proposed hierarchical segregation index (HSI). **(A)** Step 1: Sparse dictionary learning of individual resting state fMRI is performed within the SPARK framework to estimation subject-specific dictionary of functional networks and sparse coefficient matrix. Step 2. Voxel-level (*k*_*v*(*i*)_) and region-level (*k*_*r*(*i*)_) *k*-hubness for each voxel *i* are estimated from the individual sparse coefficient matrix (networks-by-voxels) using SPARK. Step 3. The hierarchical segregation index (HSI) is defined and computed for a voxel *i* as the ratio between *k*-hubness values estimated at the region-level and at the voxel-level. In this toy example, voxel 4 that has the highest number of network overlaps exhibits the lowest HSI value (i.e., more integration or less segregation within that region). *k*_*v*_ highlights voxel-level hubs participating in more than one functional network. **(B)** A nine-region-of-interest (ROI) atlas was used to estimate *k*-hubness at the region level modifying the Yeo-7 functional networks (29). The subcortical networks were defined using automated anatomical labeling (AAL) template (30): the subcortical network region and the cerebellum network region. The original Yeo-7 functional limbic network was modified to add the hippocampi and amygdalae from the AAL template. **(C)** The mean and standard deviation of regional *k*_*r*_ within the nine pre-defined regions of interest. **(D)** Group-average *k*-hubness map was obtained by averaging individual *k*-hubness maps at each vigilance state: well-rested resting state (WR), N2 sleep and N3 sleep stages. **(E)** Differences in *k*-hubness maps between states. Two-sample permutation tests (10,000 permutations) were used to reduce the bias induced by sample size difference between states. *p*_*F DR*_ *<* 0.05 (*SI Appendix, Table S1*).

## Results

We recruited 20 volunteers according to our inclusion criteria described elsewhere (12, 25, 26): participants were healthy and good sleepers aged between 18 to 30 years. We obtained and analyzed simultaneous high-density EEG-fMRI data, from the 5-minute resting state scans following a normal sleep night and the 60-minute scans during a one-hour nap inside the scanner after a whole night SD (**Fig. 1A**). Participants also performed three cognitive tasks (ANT: Attention network task (22) for testing executive attention, N-back (23)for working memory, and MCT: Mackworth clock test (24)for psychomotor vigilance) during WR as well as before and after the recovery nap following SD. Simultaneous EEG was acquired with an MR-compatible 256-channel system (Magstim EGI, Eugene, OR, USA) at 1,000 Hz. MR gradient and ballistocardiogram artefacts were then removed from EEG signals following the procedure explained in (25). Sleep stages were marked by four experts: NEC, AAP, CA and BF, for the 60-minute EEG data acquired during the nap after SD. During the one-hour nap, the total duration of N2 was 36 *±* 13.4 minutes (mean and standard deviation across subjects) and 10.8 *±* 8.8 minutes for N3 (**Fig. 1B**, *SI Appendix*). After identifying sleep states during the nap using EEG, we focused on analyzing the corresponding fMRI data at each state to estimate the functional network organizations in the whole brain, when compared to WR.

To ensure that we analyzed and compared the same number of frames per state, fMRI segments in N2 or N3 stages lasting for *>* 5 min were selected and trimmed to 5 min. We collected all 5-minute consecutive segments of N2 and N3 data. Time points with excessive motion (frame displacement *>* 0.5*mm*) were scrubbed (27). Several fMRI segments were excluded when more than 35% of time points showed excessive motion. Then, we focused on studying the first sleep cycle during the one-hour recovery nap that is suggested to capture the strongest sleep pressure after SD (28). For each subject, we selected the first 5-minute segment of N2 sleep followed by a 5-minute N3 data. In case we could not find a 5-minute N3 data immediately followed by a 5-minute segment of N2 sleep, the next available 5-minute N3 data during the one-hour scan was selected.

As a result, we considered the following clean and dimension-matched datasets for further analysis: 5-minute resting-state fMRI segments obtained from 14 subjects during WR, 18 subjects during N2, and 12 subjects during N3. The SPARK method (18) was applied to the selected fMRI datasets to estimate the functional hub organizations per state (WR, N2 or N3) for each subject. To avoid potential bias induced by differences in sample size between states, permutation tests were performed when comparing our hubness measures between states. This allowed studying hub organizations across brain states without a bias in data dimensions and focusing on the earliest measurable effects of short nap after the total SD on cognitive recovery.

### Functional network hubs reconfigure during NREM sleep

We first assessed if the global network scale *N*, i.e., the total number of networks estimated for each fMRI run, was preserved across brain states within individuals (31, 32). To estimate *N* from individual data, we applied SPARK method (18, 19) to decompose the whole brain data into several atoms, each of them being a network (21). As expected, the estimated *N* in individuals was preserved across brain states: 10.7*±* 3.4 (mean and standard deviation) for WR, 11.6 *±* 2.6 for N2 and 11.6*±*2.4 for N3 (**Fig. 1C**) after the manual removal of physiological noisy atoms. (See *SI Appendix* for details to compare *N* before and after denoising.) In the subsequent analyses, we tested our hypothesis that topological patterns of network integration is altered from WR to NREM sleep (N2 and N3), even if the global network scale *N* was preserved within individuals across different states. For each state per subject, in addition to the estimation of voxel level *k*-hubness using SPARK (*k*_*v*_: the number of networks overlapping in this voxel, usually ranging from 1 to 10 networks), we proposed a new method to estimate *k*-hubness at the region level (*k*_*r*_: the number of networks overlapping in a region of interest that involves the voxel) (**Fig. 2A, Steps 1 and 2**). To estimate *k*_*r*_, we considered nine larger networks as regions (*r* = 1, …, 9), defined using a modified version of Yeo-7 functional atlas (29) combined with two additional subcortical networks (**Fig. 2B**). As a result, we found that *k*_*r*_ increased in the visual network and decreased in the executive control (frontoparietal) network from the WR to N2 sleep state (**Fig. 2C**, Bonferroni corrected *p <* 0.05).

We then assessed how patterns of *k*-hubness estimated at the voxel level (*k*_*v*_) varied across vigilance states, as a measure of integration/segregation between networks. During WR condition, the group average *k*_*v*_ over subjects (⟨*k*_*v*_⟩_*group*_) exhibited high hubness in the default mode, frontoparietal, and visual areas (**Fig. 2D**), in agreement with our previous work estimating *k*_*v*_ (18, 19). Importantly, these patterns were found to be altered during the recovery nap following whole night SD (two-sample permutation test, FDR corrected p-value *p*_*F DR*_ *<* 0.05, **Fig. 2E**). When comparing N2 to WR states, we found decreases in *k*_*v*_ mainly within the posteior ventromedial prefrontal cortex (vmPFC) and bilateral temporo-mesial regions. In addition, we found that *k*_*v*_ then increased within some of these regions from N2 to N3 states (*p*_*F DR*_ *<* 0.05; N3 *≈* WR *>* N2), indicating sleep stage-specific changes in network overlaps. See *SI Appendix, Table S1* and *Fig. S2* for details of all voxel clusters in the thresholded *t*-score maps.

### Hierarchical segregation of brain networks during different vigilance states

The observation of distinct patterns of network integration/segregation across the region-level and voxel-level raises a question about how the networks overlapping within a region relate to the networks overlapping within a specific voxel. Studying the relative degree of network integration at a higher level (regions) with respect to those at the lower level (voxels) can provide insights into hierarchical network segregation, in a complementary manner to the metric FCR measuring network integration/segregation using mutual information within a Bayesian framework (12, 16, 17). To address this, we extended the ability of SPARK to estimate *k*-hubness at two hierarchical spatial levels, at the voxel level and at a regional level (**Fig. 2A**). **Step 1**. SPARK provides an estimation of the sparse spatial overlap between *N* functional networks at the voxel level from individual resting-state fMRI. **Step 2**. We can obtain a network (*N* : the number of whole brain networks) by voxel (the number of voxels in a region *r*) feature space for a pre-defined large region *r*, in which we estimated networks overlap. **Step 3**. We used this framework to define a new measure of network segregation within our proposed datadriven voxel-to-region hierarchy. To assess if a large functional brain system (region *r*, black dotted box in **Step 3, Fig. 2A**) was functionally segregated into several SPARK-derived networks, we define the *HSI*_*v*(*i*)_ for each voxel *i* using the ratio of region-level *k*-hubness for a region *r* to which the voxel *i* belongs (*k*_*r*(*i*)_), to voxel-level *k*-hubness (*k*_*v*(*i*)_).

This proposed HSI is equal or larger than 1, since the large-scale network in which a voxel belongs to, should involve an equal or larger number of networks than the number of subnetworks involved in this specific voxel. Therefore, a high HSI value reflects that the number of networks involved in a region *r*, to which the voxel *i* belongs to, is larger than the number of subnetworks involved in this voxel. This also reflects that this voxel may be involved in several subnetworks (*k*_*v*(*i*)_), but it belongs to a larger system that involves more subnetworks (*k*_*r*(*i*)_), whereas functional integration between those sub-networks is low. Consequently, a high HSI value suggests more segregation between subnetworks involved in the region *r*, and therefore within-region inhomogeneity. On the other hand, a low HSI value reflects that this voxel *i* belongs to a system (the pre-defined region *r*) that involves a similar number of networks than the value *k*_*v*(*i*)_. Therefore, a low HSI suggests within-region homogeneity, reflecting large integration between subnetworks involved in the region *r*.

Among them, occipital areas involving the visual network exhibited increased HSI values during N2 compared to WR, followed by decreased HSI values during N3 compared to N2, ultimately returning to the levels comparable to those observed during WR (**Fig. 3B**). The posterior vmPFC showed decreased HSI values during N2 compared to WR, with values returning to wakefulness levels during N3 compared to N2. Given the discrepancy between region-level k-hubness (**Fig. 2C**) and voxel-level k-hubness (**Fig. 2D**) in the visual cortex across vigilance states, the HSI-based findings provide new insights into how large-scale networks segregates into smaller subnetworks. In contrast, parts of the posterior cingulate cortex (PCC) within the default mode network showed increased HSI during N2 compared to WR, with no significant further changes between N2 and N3. Nevertheless, HSI in the PCC remained modestly elevated during N3 relative to WR, indicating that the N2-related increase in HSI persisted into deeper sleep. This pattern differs from that observed in the visual network, where substantially larger HSI changes occurred between N2 and N3. More detailed analysis (*SI Appendix, Table S2*) additionally suggested decreased network segregation in the temporo-mesial structures, including bilateral hippocampi, during N2 compared to WR.

**Fig. 3.**
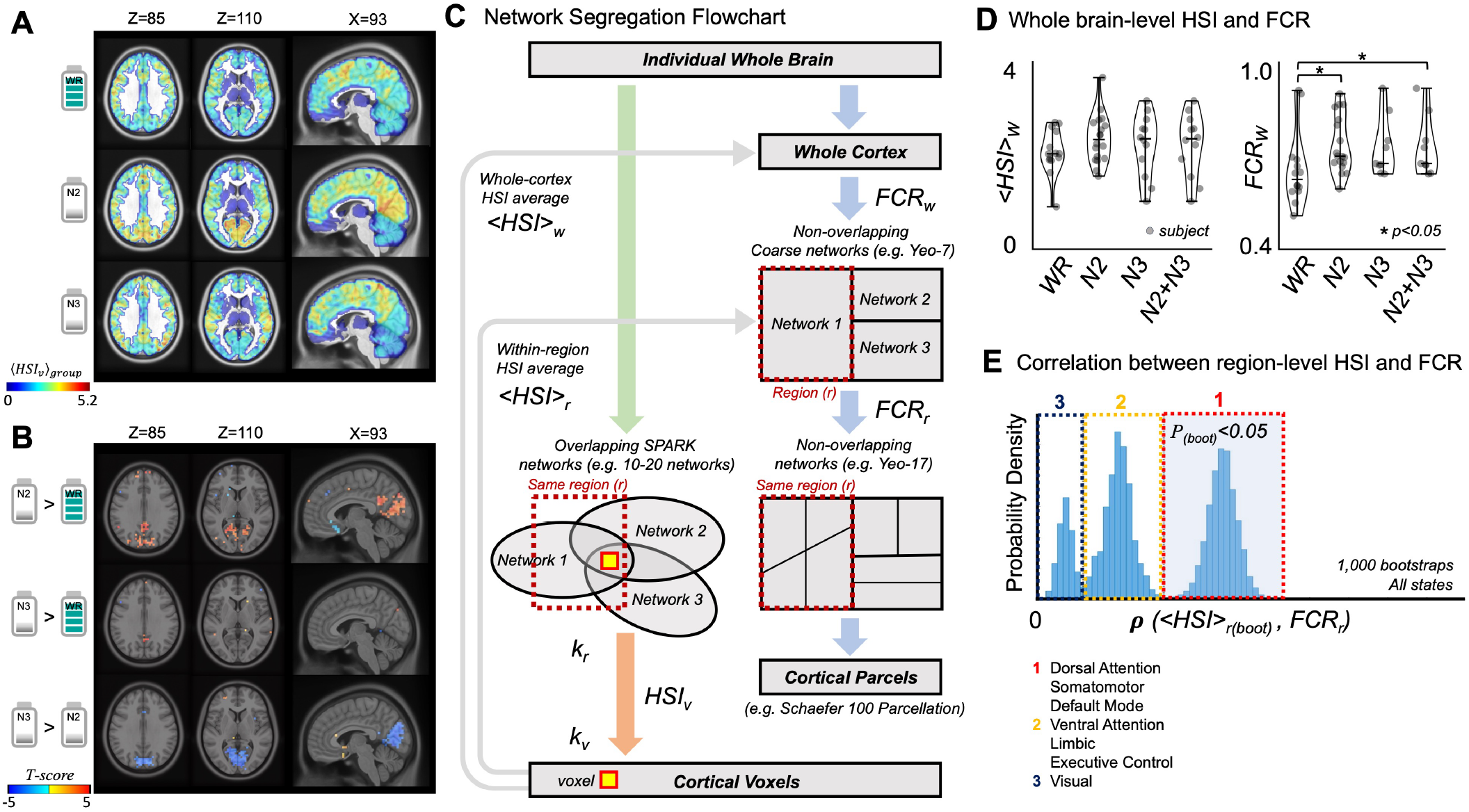
Altered patterns of functional network segregation across different vigilance states. Voxel-level HSI can identify subtle changes of stage-specific patterns of network segregation during sleep. **(A)** Group average HSI maps estimated from wakeful resting state (WR), NREM2 sleep (N2) and NREM3 sleep (N3). Voxel-level HSI values (*HSI*_*v*_) were averaged across subjects. **(B)** t-statistics map comparing HSI between states: N2 *>* WR (top), N3 *>* WR (middle), and N3 *>* N2 (bottom). Two-sample permutation tests, 10,000 permutations, *p*_*F DR*_ *<* 0.05 (*SI Appendix, Table S2*). **(C)** Segregation flowchart. For each individual resting-state fMRI, SPARK was applied to estimate overlapping networks (green arrow) and the corresponding HSI at the voxel level (orange arrow). *⟨HSI⟩*_*w*_ and *⟨HSI⟩*_*r*_ is the average of voxel-level HSI values (*HSI*_*v*_) estimated over the whole cerebral cortex (*w*) and the average within each region (*r*). A region *r* is defined using a non-overlapping coarser network (red dotted boxes). Since subcortical and mesial temporal structures were not included in Schaefer100 parcellation considered for *FCR*_*w*_ estimation, we also excluded them when estimating whole cortex averaged *⟨HSI⟩*_*w*_. *FCR*_*w*_ and *FCR*_*r*_ quantify hierarchical segregation of the entire cortex (subscript *w*) into the non-overlapping Yeo-7 networks and each of Yeo-7 network (considered as a region *r*) into its sub-networks (blue arrows). **(D)** Network segregation estimated in the whole brain cortex using *⟨HSI⟩*_*w*_ (left) and *FCR*_*w*_ (right). Wilcoxon rank sum test, *p <* 0.05. **(E)** Distribution of the correlation coefficients (*ρ*) between *⟨HSI⟩*_*r*(*boot*)_ and *FCR*_*r*_ obtained using a bootstrap resampling approach for the reliable estimation of *⟨HSI⟩*_*r*_. To obtain robust mean values of *HSI*_*v*_ within one brain region and to mitigate information loss resulting from averaging *HSI*_*v*_ across an extended brain area, *⟨HSI⟩* _*r*(*boot*)_ values were estimated by a spatial bootstrap approach for each region *r*. Three patterns of level of correlation were found from this histogram. Groups were defined using two local minima *ρ* = 0.02 and 0.23.

### Voxel-level HSI can identify subtle changes of stage-specific patterns of network segregation within regions: Comparisons to FCR

Developed within the framework of a hierarchical Bayesian model of network integration, the Functional Clustering Ratio (FCR) is a complementary metric defined as the ratio of the integration within subsystems and the integration between subsystems. Originally proposed by Marrelec et al. (17) and adapted by Boly et al.(16), FCR was used for studying network segregation during NREM sleep in our previous work (12). In the present work, we focus on studying different patterns of network segregation using the HSI metric, specific to N2 and N3 sleep stages during the recovery nap following SD, in comparison to WR. The HSI and FCR methods are relying on different assumptions about network reorganizations and hierarchy (**Fig. 3C**). First, HSI allows generating a whole brain map of segregation quantified at the voxel level (*HSI*_*v*_), whereas FCR quantifies network segregation as a single value for a large network of interest in an association with its sub-networks within a hierarchical model (*FCR*_*w*_ for the whole cortex or *FCR*_*r*_ for a region *r*). Second, HSI considers a data-driven network hierarchy based on overlapping functional networks estimated using SPARK, and the anatomical boundaries of these subnetworks are not limited by the selected regions of interest (*r*). On the other hand, FCR considers network hierarchy of non-overlapping functional networks (e.g. Yeo 7 and 17 networks as in (12)), in which the anatomical boundaries of sub-networks are directly limited by their high-order network.

Here, we assume that the estimation of voxel-level HSI using data-driven overlapping network structures can provide information complementary to FCR that is measured at the regional level assuming a pre-defined hierarchy of non-overlapping networks. The proposed analysis of *HSI*_*v*_ can address within-region heterogeneity of network segregation that cannot be addressed using FCR. To assess this assumption, we compared HSI and FCR from the same datasets. We performed two analysis comparing the behavior of HSI and FCR metrics, first at the whole cortex level and then at the regional level (**Fig. 3C**).

### Comparison of HSI and FCR at the whole cortex level

At the whole cortex level, the *FCR*_*w*_ was compared to the mean HSI over the voxels in the whole cortex (*⟨HSI⟩*_*w*_) (**Fig. 3D**). The overall trend of increase in network segregation from WR to NREM sleep was found using both ⟨*HSI*⟩_*w*_ and *FCR*_*w*_ values. The estimated ⟨*HSI*⟩_*w*_ was 2.08 *±* 0.47 during WR, 2.36 *±*0.52 during N2 sleep, 2.23*±* 0.67 during N3 sleep, and 2.31*±* 0.58 when combining N2 and N3 sleep states. The *FCR*_*w*_ increased from 0.66 *±*0.12 in WR to 0.74 *±*0.09 during N2 (Wilcoxon rank sum test, *p <* 0.05), 0.73 *±*0.1 during N3 sleep, and 0.73*±*, 0.1 when combined N2 and N3 (*p <* 0.01). Network segregation estimated using ⟨*HSI*⟩_*w*_ and *FCR*_*w*_ therefore exhibited similar trends when comparing vigilance states, but only *FCR*_*w*_ demonstrated a significant increase in segregation from WR to N2. This might be explained by the appropriate hierarchical pooling of information theory measures along a predefined hierarchy, resulting in reliable measures of network segregation at the whole brain or network levels. On the other hand, *FCR*_*w*_ showed no difference between N2 and N3 states. In addition, the expected subsequent decreases in network segregation from N2 to N3 even during NREM sleep, as suggested by our analysis of HSI at the voxel-level (**Fig. 3B**), was not retrieved at this regional spatial scale using neither ⟨*HSI*⟩ _*w*_ nor *FCR*_*w*_.

### Comparison of HSI and FCR at the regional level

At the regional level, after pooling results from all states (WR, N2, N3), *FCR*_*r*_ was compared to the mean HSI within each corresponding Yeo7 cortical region (⟨*HSI*⟩_*r*_) (**Fig. 3E**). To do so, we first averaged the HSI values within each of the Schaefer100 parcels, resulting a single HSI value for each of the 100 parcels. Second, a bootstrap approach was employed to obtain robust statistics for the association between the ⟨*HSI*⟩_*r*_ and *FCR*_*r*_, to prevent a potential bias induced by within-region inhomogeneity when averaging HSI values over large regions. For each region *r* that involves *m*_*r*_ parcels, a bootstrap replication was generated by resampling a new set of *m*_*r*_ parcels with replacement, within Schaefer100 parcels of the regions *r*. Then, ⟨*HSI*⟩_*r*(*boot*)_ for this bootstrap sample was computed by averaging the values in the resampled set. A total of 1,000 bootstrap samples were considered. Third, the correlation coefficient (*ρ*_(*boot*)_) between *FCR*_*r*_ for all subjects and vigilance states with each bootstrap surrogate ⟨*HSI*⟩_*r*(*boot*)_ and the corresponding *p*value (*p*_(*boot*)_) were calculated (*SI Appendix Fig. S3*).

After pooling results from the 7 Yeo networks, the resulting distribution of correlation values *ρ*_(*boot*)_ revealed the presence of three groups of regions, suggesting different patterns of HSI-FCR correspondence over the cerebral cortex. Specifically, Group 1 that exhibits significant correlation between ⟨*HSI*⟩_*r* (*boot*)_ and *FCR*_*r*_ (*ρ*_(*boot*)_ = 0.38 *±* 0.04, *p*_(*boot*)_ *<* 0.05) mainly included regions of the dorsal attention, somatomotor, and default mode networks. Group 2 reporting subtle non significant correlations (*ρ*_(*boot*)_ = 0.11*±*0.04) included the ventral attention, limbic and executive control networks. Group 3 reporting no correlation (*ρ*_(*boot*)_ = −0.03 *±*0.02) included the visual network. Among the three networks of Group 1, the dorsal attention network exhibited the largest correlation between the two measures, suggesting similar patterns of segregation assessed with those complementary metrics (*SI Appendix Fig. S3*).

Together, our results demonstrate that the within-region heterogeneity of network segregation across NREM sleep stages (N2 versus N3) can be retrieved at the voxel level and that the proposed voxel-level analysis of HSI was more sensitive to local changes in network segregation than using either HSI or FCR estimated at the whole-brain or regional level. We also found that the correspondence between the two proposed metrics of segregation (HSI and FCR) was inhomogeneous over the cerebral cortex, highlighting the largest HSI- FCR correlation in the dorsal attention, somatomotor, and default mode networks and no correlation in the visual network.These results aligned with the voxel-level HSI analysis, where we found increased HSI in the PCC, a key hub of the default mode network, during N2 relative to WR (N2 *>* WR). From N2 to N3, no significant difference was observed in the PCC (N3*≈* N2), although HSI remained modestly elevated in N3 relative to WR (**Fig. 3B**). In contrast, our voxel-level analysis revealed N2- and N3-specific network reorganization characterized by (i) an increase in HSI in the visual network from WR to N2 followed by a decrease from N2 to N3 (N2 *>* N3 *≈* WR), and (ii) a decrease in HSI in the posterior vmPFC from WR to N2 followed by an increase from N2 to N3 (N2 *<* N3 *≈* WR) (**Fig. 3B**).

### Impacts of network segregation during NREM sleep on cognitive recovery from SD

We then investigated whether the observed functional network segregation during NREM sleep was associated with cognitive recovery from SD. As previously reported by Cross et al. using FCR for the same dataset, the cognitive performance after total SD significantly dropped when compared to WR (12). After taking the nap, the cognitive performances then significantly increased when compared to SD, but still remained lower than when compared to WR (12). In this study, we focused on studying the role of NREM sleep stages N2 and N3 in the recovery of cognitive task performances from SD. To address this, we evaluated the association between the overall level of network segregation within each region ⟨*HSI*⟩_*r*_ during N2 and N3 sleep stages and the improvement of cognitive performances after taking the nap, when compared to the SD state before recovery nap (PRN*>*SD, **Fig. 4**). Since SPARK method provides an absolute measure of hubness as integers ranging from 0 to less than *N* networks, we can consider HSI values during NREM sleep without performing any normalization to a baseline value. To assess individual cognitive performance of N-back, MCT and ANT tasks during either PRN or SD states, we measured the task reaction time (RT) in milliseconds (ms) as well as the task accuracy, i.e. the percentage (%) of correct responses.

**Fig. 4.**
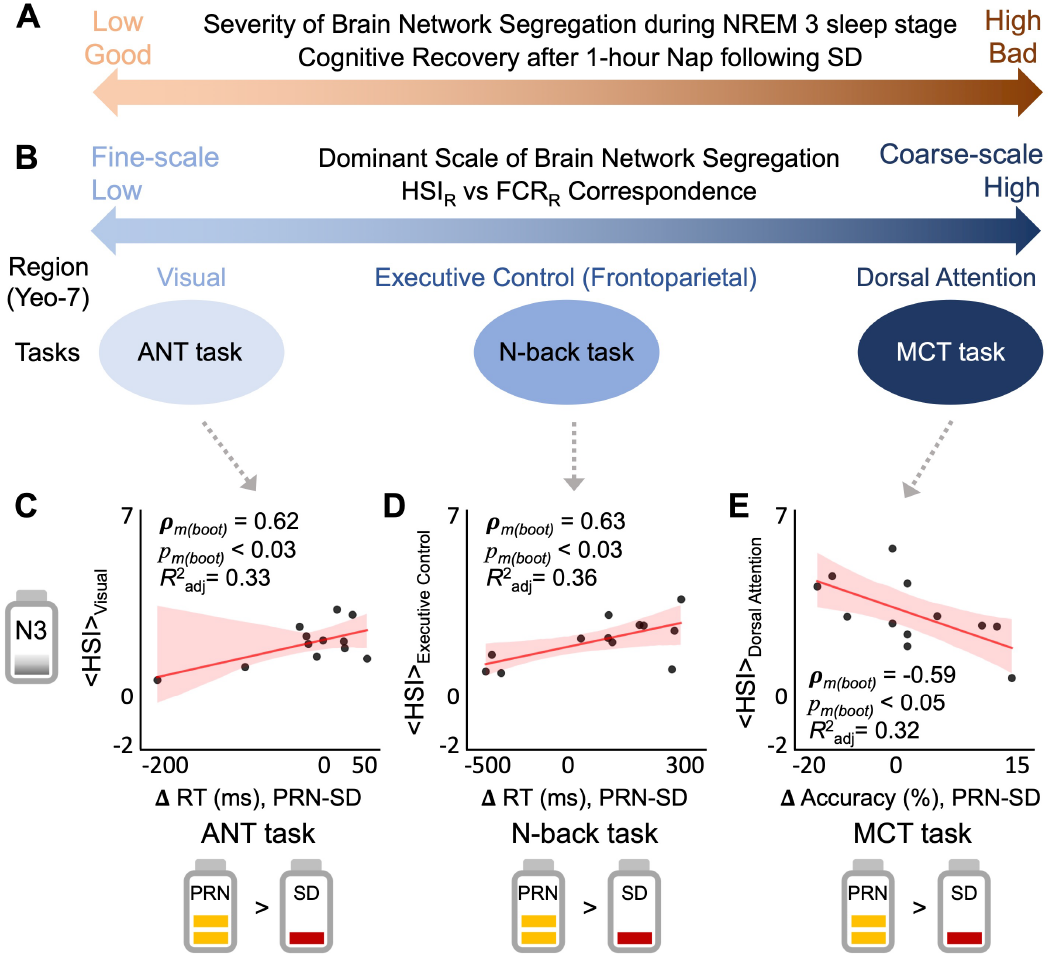
The severity and spatial scale of brain network segregation during N3 sleep stage impacts individual’s ability to recover cognitive task performances after 1-hour nap following 24-hour sleep deprivation (SD). **(A)** Individuals exhibiting more brain network segregation in NREM 3 sleep stage during 1-hour nap following sleep deprivation showed worse recovery of cognitive functions. **(B)** The level of spatial resolution of network segregation differed between brain regions and was associated with the recovery of distinct cognitive tasks. We tested all possible region–task pairs and present only the associations that survived statistical testing. The correspondence between ⟨*HSI*⟩_*r*_ and *FCR*_*r*_ reflects at which spatial resolution brain network segregation was dominantly observed (region-level or voxel-level). **(C-E)** Scatter plots supporting the summary in **(A-B)**. y-axis denotes the HSI estimated at the region level (⟨*HSI*⟩_*r*_), where the regions (*r*) are the visual **(C)**, executive control **(D)**, and dorsal attention **(E)** networks defined using the Yeo-7 functional parcellation. x-axis denotes the changes in task performance measured before and after the nap (post-recovery nap vs before-recovery nap; PRN-SD), when considering ANT **(C)**, N-back **(D)**, and MCT **(E)** tasks respectively. Black data points are individual subjects. Red regression lines represents the best fit line through the data points. Red shaded area around the regression line indicates a 95% confidence interval. Adjusted coefficient of determination 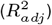 was estimated using the fitted linear regression model. *ρ*_*m*(*boot*)_ and *p*_*m*(*boot*)_ denotes the median of *ρ*_(*boot*)_ of p-value *p*_(*boot*)_ across 1,000 bootstrap samples, using our spatial bootstrap strategy to achieve a robust estimation of the neurobehavioral association. N3, NREM 3 sleep stage; PRN, post recovery nap; SD, sleep deprivation.

As a result of testing all possible region-task pairs, we found that more network segregation during N3 sleep in the recovery nap (e.g. high *HSI*_*r*_ values) was associated to poorer post-nap recovery of cognitive task performances: ANT task (executive attention), N-back task (working memory), and MCT task (psychomotor vigilance) (**Fig. 4A**). Network segregation commonly found during N2 sleep was not associated with any change task performances. Moreover, we found that network segregation in different brain regions impact the recovery of distinct cognitive tasks. Specifically, the regional average of network segregation in the visual network was associated with the recovery of ANT performance after the nap (**Fig. 4C**, reporting reaction time). The observed network segregation in the executive control region was associated with the recovery of N-back task performance (**Fig. 4D**, reporting reaction time). Finally, network segregation of dorsal attention region during N3 sleep was associated with the recovery of MCT performances (**Fig. 4E**, reporting task accuracy). We did not find any association in other regions with any tasks. Interestingly, these regions were also involved in different groups when comparing HSI and FCR metric: the dorsal attention region exhibited the strongest HSI-FCR correlation (Group 1). The executive control region exhibited subtle HSI-FCR correlation (Group 2), and the visual network exhibited no correlation (Group 3) (**Fig. 3E**). On the other hand, when considering comparison at the voxel level, it was the visual network that exhibited the most significant changes from three states (N2 *>* WR *≈* N3). These results together suggest that dominant spatial scale (resolution) of network segregation differs between brain regions and impacts the recovery of distinct cognitive tasks (**Fig. 4B**).

In addition, we compared the recovery of task performances (PRN*>*SD) to the difference in HSI between NREM sleep stages and the WR state as a baseline (Δ⟨*HSI*⟩_*r*_, N2 or N3*>*WR). In these comparisons, we confirmed our finding that more segregation during N3 sleep was associated with poorer recovery of task performances. However, only the MCT task performances (PRN*>*SD) showed such association with HSI during N3 when compared to WR (Δ⟨*HSI*⟩ _*r*_, N3*>*WR) both in the dorsal attention network regions (Accuracy; *p*_*m*(*boot*)_ *<* 0.05, *ρ*_*m*(*boot*)_ = *−*0.753) and the visual network (Accuracy; *p*_*m*(*boot*)_ *<* 0.05, *ρ*_*m*(*boot*)_ = *−*0.708).

## Discussion

State-dependent variation of brain activity in hubs associated with overlapping network structures in the whole brain have not been previously described at the voxel level during NREM sleep. Here, we developed a new metric, the hierarchical segregation index (HSI) to quantify the degree of segregation between and within functional brain networks, offering voxel-level mapping of these properties. To do so, we applied the SPARK method to estimate *k*-hubness, the number of overlapping networks in each voxel, using sparse dictionary learning for individual resting state fMRI (18, 19). In this work, we extended the ability of SPARK to provide the estimation of *k*-hubness at the region level and at the voxel level at the same time for a pre-defined large-scale region (**Fig. 2C**). This allowed us to define the HSI for each voxel as the ratio of *k*-hubness assessed at the region-level (*k*_*r*_) and at the voxel-level (*k*_*v*_), where the voxel belongs to this region *r*.

Using the proposed metric HSI, we found that network segregation increased in the PCC, a core region of the default mode network, during NREM sleep following SD, when compared to well-rested wakefulness (**Fig. 3B**). This is in agreement with previous findings using FCR (12, 16), which focused on the contrast between WR and NREM sleep, but combining N2 and N3 sleep stages in the same measure. In addition, we found that segregation in the visual network and the posterior vmPFC exhibit N2 and N3 sleep specific voxel-level patterns. More segregation of brain networks during N3 sleep was associated with worse retrieval of cognitive functions after the nap, whereas we found no relationship between network segregation during N2 sleep and change in task performance.

By evaluating the correspondence between HSI to FCR (12, 16) measured across different spatial scales (voxel, region, and within the entire cortex), our study revealed that brain regions exhibiting different dominant spatial scale of segregation impacted the recovery of different cognitive tasks. Visual network showed the largest N2 and N3 sleep specific changes in HSI at the voxel level (**Fig. 3B**), while exhibiting no correspondence between HSI and FCR at the region level (**Fig. 3E**). Moreover, segregation within the visual network during N3 was also associated with worse recovery of ANT performance after the nap. On the other hand, the dorsal attention network exhibited the strongest correspondence between HSI and FCR at the region level, and more segregation within this network during N3 was associated with worse recovery of MCT performances. Importantly, by focusing on N2 versus N3-specific patterns of network alterations rather than treating NREM sleep as a homogeneous state, this work reveals a distinct role of N3 sleep in cognitive recovery, suggesting a modulatory influence of local network integration operating at a finer scale than that captured by traditional region-level analyses.

Together, our study demonstrates that the proposed analysis of voxel-level HSI can provide information complementary to the conventional region-level measure of network segregation, as provided by FCR metric. Our analysis reports key insights into within-region variation that cannot be addressed at the regional level, suggesting a mechanistic understanding of how NREM sleep replenishes cognition after sleep deprivation.

### Preserved global network scale across vigilance states

First, we observed vigilance state-specific reorganization of functional hubs occurring during NREM sleep, whereas the global network scale, i.e. the total number *N* of networks estimated in each state for each subject, was preserved (**Fig. 1C**). Using the automatic estimation of global network scale estimated with the minimum description length criteria within the SPARK framework (18, 19), the estimated global network scales in the present datasets were found similar during WR, N2 and N3 states. This result is in line with previous findings of the preservation of global network scales from healthy young adults across different arousal levels during resting state, using SPARK at the region level (20), and from comatose patients in the absence of consciousness using graph theory at the region level (31). It suggests that brain activity preserve a specific amount of complexity at the level of whole brain networks within individuals, while local functional reorganizations of brain network interaction were specific to each vigilance state.

### Altered integration during N2 and N3 sleep stages at the region level

We reported decreased region-level *k*-hubness (*k*_*r*_) in the frontoparietal (executive control) regions from WR to N2 sleep states and increased *k*_*r*_ from N2 to N3 sleep (**Fig. 2C**). Using independent component analysis of resting state fMRI data during awake states with and without SD, Kaufmann et al found SD-induced alterations of brain functional connectivity, mainly between the default mode, dorsal attention, frontal and visual networks (33). Using hierarchical cluster analysis of frontoparieal functional connectivity estimated from resting state fMRI, Spoormaker et al. demonstrated a transition from a globally integrated functional brain network in WR to segregated networks during sleep (13). They suggested a key role of frontoparietal hubs to support these reorganizations. Using SPARK, the values of *k*-hubness can be interpreted as the level of functional complexity in a region *r*, indicating how many local and long-range functional processes are concurrently involved in *r* (18). In this context, the decreased *k*-hubness during N2 reflects reduction of long-range functional connectivity of the frontoparietal regions, whereas the increased *k*-hubness during N3 reflects enhanced frontoparietal local connectivity.

In the visual cortex, regional-level *k*-hubness (*k*_*r*_) increased from WR to N2 and then decreased from N2 to N3 (**Fig. 2C**). Note that we used permutation tests to avoid potential bias induced by differences in sample size between states. These results suggest that state-specific network reorganization involve mixed patterns of network integration associated with local and long-range connectivity in visual hubs, and that the direction of sleep-dependent changes in network integration differ between the visual and frontoparietal regions. Previously, seed-based analysis using seeds in the calcarine gyri showed increased local connectivity (segregation) in the visual cortex from WR to N2 and further to N3 (13), in agreement with our findings on reduced visual network segregation (increased integration) during N3. They also reported decreased thalamus-visual (long-range) functional connectivity from WR to N2 followed by a slight increase from N2 to N3 (13). Similarly, a low arousal resting state assessed using decreased pupil area during wakefulness was associated with increased *k*-hubness in the visual network and decreased *k*-hubness in the frontoparietal network (20).

### Altered integration during N2 and N3 sleep stages at the voxel level

The investigation of *k*-hubness estimated at the voxel level (*k*_*v*_) further revealed the largest reduction of *k*_*v*_ from WR to N2 in the posterior vmPFC and the mesiotemporal structures including bilateral hippocampi (**Fig. 2E**, *SI Appendix Table S1*). Given the proposed role of sleep spindles in memory consolidation during N2 (34) and hippocampal-vmPFC connectivity (35), future studies investigating the relationship between the decreased *k*-hubness in mesiotemporal structures observed in the present work and sleep spindles may help clarify whether these network changes reflect spindle-related hippocampal-cortical interactions during N2.

### Increased network segregation in the PCC during NREM sleep after SD compared to well-rested wakefulness

Using HSI, we first found an overall increase in network segregation mainly in the PCC during NREM sleep when compared to WR, whereas we found no further changes when comparing N2 and N3 states (**Fig. 3B**, *SI Appendix Table S2*). It suggests that increased network segregation in these regions could be a marker of loss of consciousness during sleep when compared to WR, similar to the findings reported using FCR metric on task fMRI when subjects performed the three cognitive tasks (after regressing out the effect of the task) from the same dataset (12). Indeed, in our previous study, Cross et al. reported increased network segregation at the whole cortex level (increased *FCR*_*w*_) during NREM sleep when compared to WR and SD states, followed by decreased *FCR*_*w*_ during tasks after the nap (WR*<*PRN*<*SD*<*NREM) (12). At the region level, we reported increases in *FCR*_*r*_ during NREM sleep in the default mode, somatomotor, ventral attention and executive control networks, when compared to WR (12). We also found that FCR further increased during the entire 1-hour nap containing NREM sleep after SD, suggesting that the altered balance of within-networks and between-networks integration could indeed be a marker of consciousness (12). In line with these findings, our present work found strong correspondence between HSI and FCR metrics estimated from the same data within the default mode, dorsal attention, and somatomotor regions (**Fig. 3E**). Together, the results provide converging evidence that the increased hierarchical network segregation in the PCC to be a hallmark of consciousness during NREM sleep.

### Different patterns of voxel-level network segregation in the visual network and posterior vmPFC during N2 and N3 sleep stages

Using HSI metric at the voxel level, we found increased network segregation in the visual network during N2 sleep stage when compared to WR, which then decreased from N2 to N3 states (N2*>*WR *≈*N3, **Fig. 3B**). On the other hand, in the posterior vmPFC, we found decreased network segregation during N2 when compared to WR, which then increased from N2 to N3 stages (N2*<*WR *≈*N3). Using hierarchical clustering and a Bayesian framework (17), Boly et al. reported increased segregation mainly in the visual, salience, executive control and dorsal attention networks during NREM sleep when compared to WR (16). However, they did not study N2 or N3 sleep specific patterns of network segregation. Our work demonstrate that N2 or N3 sleep stages exhibit unique patterns of network segregation, in addition to the overall network reorganization during NREM sleep in comparison to WR. Specifically, a pattern found when comparing N2 sleep to WR was reversed when comparing N3 to N2 sleep stages. Moreover, the limbic and visual networks showed relatively low correspondence when we compared HSI with FCR at the region level (**Fig. 4**). These findings together suggest that the local voxel-level change is a dominant feature of brain network reorganization in these regions, contributing to the different patterns found during N2 and N3 sleep stages, whereas those effect might be diluted when analyzing those data at the regional level. Our proposed analytic framework and the development of our new metric HSI, can provide key insights into within-region variation that cannot be addressed at the region level.

Previous fMRI studies reported reduced vmPFC activity during fear memory consolidation tasks and decreased resting-state functional connectivity linked to risk-taking behavior after whole-night sleep deprivation (36, 37). Although these findings suggest that the vmPFC is vulnerable to sleep disruption, its activity during subsequent sleep has not been investigated. Our findings address this gap by demonstrating N2 and N3-specific voxel-level patterns of vmPFC network segregation after sleep deprivation.

### More segregation of brain networks during N3 sleep is associated to worse retrieval of cognition from sleep deprivation

It has been suggested to be critical for cognitive function to maintain a balance between integration and segregation of information flow between distributed brain networks (38, 39). At the whole cortex level, in Cross et al. we reported a bidirectional effect of SD-induced network segregation using the same dataset. We found that individuals exhibiting more segregation (increased *FCR*_*w*_) during task fMRI performed in SD condition (WR*<*SD) showed more decline in task performances, while individuals exhibiting more segregation during task fMRI in SD condition when compared to PRN condition (SD*>*PRN) showed more improvement of task performances after the nap (12). However, we did not consider spatial variability of such neuro-behavioral associations and combined behavioral metrics from three cognitive tasks using principal component, without considering potential differential impacts on these cognitive tasks separately. Importantly, the impact of network segregation during NREM sleep on cognitive recovery was unexplored in this previous work. Our present work fills this gap by comparing the ⟨*HSI*⟩_*r*_ estimated during NREM sleep to the changes in task performances before and after taking the recovery nap (PRN-SD, **Fig. 4**). We found that more segregation of brain networks during NREM sleep was associated with worse retrieval of working memory, executive attention, and psychomotor vigilance after the nap. Furthermore, by exploring these associations during N2 and N3 sleep separately, our analysis suggests an important role of N3 sleep, also known as slow wave sleep or deep sleep (1), modulating the maintenance of distinct cognitive functions.

Previous work suggested that N3 sleep is related to sleep homeostasis, processing of information related to the experiences of the previous awake period(4), and memory consolidation (40–42). Our results extend these findings by suggesting N3 sleep-specific roles of network reorganization in different brain regions in the recovery of working memory, executive attention, and psychomotor vigilance from SD. It should be further investigated why more integration during N3 is associated to better recovery of task performances.

### Dominant spatial scale of N3 sleep network segregation differs between brain regions and impacts the recovery of distinct cognitive tasks

By investigating the correspondence between HSI and FCR across spatial scales (**Fig. 3**) and the associations between HSI and cognitive task performances (**Fig. 4**), our results suggest an interesting relationship between the spatial resolution (scale) of network segregation, the spatial distribution (brain regions) exhibiting network segregation, and the type of cognitive task performances (brain function) that recover after 1-hour nap following SD. In our study, the visual region mainly exhibited a fine-scale network reorganization (at the voxel-level), and was associated with the recovery of executive attention function after the nap. In contrast, the dorsal attention region mainly exhibited coarse-scale network segregation (exhibiting the largest HSI/FCR correlation at the region level) and was associated with the recovery of psychomotor vigilance. The executive control network regions only exhibited subtle correspondence between HSI and FCR, and was associated with the recovery of working memory (**Fig. 4**). Importantly, N3 sleep stage, more than N2, played a key role in those relationships.

Our observations suggest that the recovery of distinct cognitive functions affected by total SD may recruit distributed brain regions and diverse hierarchical neural processes during NREM sleep. Luber et al. reported a task-specific effect of fMRI-guided repetitive TMS (rTMS) in working memory function of sleep-deprived individuals (43). They found that rTMS to left lateral occipital cortex remediated a delayed match to sample task performance (working memory) in sleep-deprived individuals, but not the performance of the psychomotor vigilance task (43). It is also suggested that some individuals are less cognitively vulnerable to SD than others (44). Individual differences in the cognitive vulnerability to SD and the ability to recover cognitive functions from SD may be associated to individual-specific spatial patterns of network reorganization during NREM sleep and the scale of network segregation (45). To the best of our knowledge, this is the first study that demonstrated different patterns of functional brain network segregation across different spatial scales (from the whole cortex to the voxel level) and across different NREM sleep stages (N2 and N3 separately), in relation to distinct cognitive functions.

### Methodological considerations and limitations

SPARK is an original sparse decomposition approach, which does not require conventional pairwise connectivity matrix and graph theory to compute hubness (46). We should interpret with caution our *k*-hubness findings, for example, the identification of *k*-hubness is not interchangeable with connector hubs in graph theory. *k*-hubness is, by definition, the number of overlapping networks in each voxel (*k*_*v*_) or region (*k*_*r*_), regardless of the size and spatial distribution of the networks. The values of *k*-hubness can be interpreted as the level of functional complexity in a voxel or a region, indicating how many local and long-range functional processes are concurrently involved in each voxel. Therefore, a large *k*- hubness value estimated from SPARK can be observed either in provincial or connector hubs defined using graph theory.

In our study, we carefully compared FCR(17) and HSI metrics of network segregation at different spatial levels, involving comparison at the whole brain and regional levels. We are reporting moderate and heterogeneous levels of concordance. At the whole brain level, *FCR*_*w*_ exhibited significant results, suggesting overall increase network segregation during N2 and N3, whereas only a trend (not significant) was found for *HSI*_*w*_. On the other hand, only *HSI* metric was able to demonstrate promising results regarding local reorganization of brain network during the recovery nap and their relationship with improvement of cognitive performance after the nap. This might be explained by the appropriate Bayesian model pooling measures of information theory over different spatial scales in FCR, resulting in reliable measurements at large spatial scales. Whereas in this study we considered a network hierarchy imposed by Yeo networks

(29) as proposed in (12) by Boly et al. estimated FCR metrics using a hierarchy of brain networks derived from the data, using independent component analysis and spatial clustering approaches (16). It would be interesting to combine the data-driven network hierarchy proposed by SPARK, with FCR estimation on similar networks than HSI, therefore fully exploiting the complementarity between the two approaches. This investigation was out of the scope of this paper.

Secondly, we did not include the subcortical regions including the basal ganglia, cerebellum, hippocampi and amygdalae in our FCR estimation, due to the mathematical constraints associated with the computation of FCR. Actually, FCR was computed from the estimation of a data covariance matrix that requires tuning a trade-off between the duration of scan and spatial scale to avoid an ill-posed estimation (17). In Cross et al., in addition to comparing Yeo-7 (upper-level) and Yeo-17 (lower-level) parcellations, we also estimated FCR for each Yeo-17 parcel in a second-level hierarchy, in which the Yeo-17 (upper-level) parcellation was compared with a smaller 57 data-driven clusters identified from 400 parcels covering the whole brain. This was only possible because FCR data were estimated from long duration task fMRI data, obtained by concatenating fMRI time-series acquired from the three cognitive tasks (after regressing out the task paradigm). To ensure that similar amounts of sleep data could be considered in such analysis, the entire duration of NREM sleep combining N2 and N3 needed to be analyzed as a single sleep state in (12).

It would be interesting to compare our results using the same second-level hierarchy, however, the present work focused on sleep-associated network alterations when compared to the short duration of resting state fMRI data (5 minutes of acquisition). Here, we analyzed only a continuous 5-minute segment of N2 and N3 sleep datasets respectively to match with the total duration of resting state data, to avoid any bias in sparse dictionary learning and network estimation procedures induced by the length of data. In addition, the present work focused on studying the first sleep cycle during the one-hour recovery nap. We first identified a set of 5-minute consecutive N2 and N3 data. Then, we selected the first 5-minute segment N2 data that was followed by a 5-minute N3 data. If a 5-minute N3 sleep segment immediately followed by a 5-minute N2 segment was not found, we selected the next available 5-minute N3 segment within the one-hour scan window. A limitation of this approach is that we cannot assume that sleep stages are the same along different cycles and the same number of sleep cycles during the 1-hour nap in our experiments.

When compared to Cross et al. (12), a smaller number of subjects was included in this study, due to our conservative data screening criteria for motion parameters, which is a limitation of this study. To this end, the proposed analysis of HSI using SPARK demonstrated the ability to capture meaningful patterns of sleep-associated network segregation using the short duration of resting state fMRI data without imposing data duration constraint as for the estimation of FCR. The analytic framework of SPARK involves a bootstrap resampling strategy to find consistent patterns of resting state networks that are reproducible across resampled data. By extending the direct outcome of SPARK analysis and defining HSI metric using k-hubness at two spatial scales, our proposed method allows to separate the effects of N2 and N3 sleep on the functional network reorganizations and hubs. In future work, it would be interesting to study dynamic evolution of networks hubs during sleep and in individuals with mental illness associated with sleep disturbance (47), taking advantage of the sensitivity and reliability of SPARK demonstrated in this and our previous work (18–20).

### Conclusions

By developing a new voxel-level measure of hierarchical network segregation for individual resting state fMRI, our study provides evidence of the reorganization of functional brain networks during a recovery nap after sleep deprivation, suggesting a role of NREM sleep in restoration of working memory, executive attention and psychomotor vigilance functions. We identified specific patterns of network segregation of the PCC in the default mode network characterizing the overall NREM sleep relative to wakefulness, therefore suggesting that network segregation could serve as a marker of consciousness. We further observed dissociable sleep-stage–dependent segregation patterns in the posterior vmPFC and visual network regions, such that segregation increased in the visual cortex and decreased in the posterior vmPFC during N2, before returning to wakefulness-like levels during N3. More integration of brain networks during N3 sleep was associated with better retrieval of task performances during a recovery nap after SD. Importantly, our results showed that dominant spatial resolutions of network segregation differed between brain regions, being associated with the recovery of distinct cognitive tasks. Our study suggests the importance of considering different characteristics of brain network reorganizations across brain regions that may occur at different spatial scales (from whole brain to networks to voxels). Moreover, the correspondence between the state-dependent changes of HSI and FCR were heterogeneous over the cerebral cortex, demonstrating the sensitivity and reliability of voxel-level HSI to provide key insights into within-region variation that cannot be addressed at the region level. Taken together, this work provides a detailed understanding of neural basis of cognitive recovery during NREM sleep in the 1-hour recovery nap following SD.

## Materials and Methods

This study has been approved by le Comité Central d’Éthique de la Recherche du ministre de la Santé et des Services Sociaux (CCER) in Montréal, Québec, Canada. The data acquisition was performed and completed in the Sleep Laboratory and in the Bioimaging Unit of PERFORM Centre, Concordia School of Health, in Concordia University, Montreal, Quebec, Canada. The same dataset was used in our previous work (12, 25, 26).

### A 3-visit study protocol

During the first night, we used a whole-night polysomnography (PSG) monitoring for participant screening purposes. Once a participant was considered eligible, a two-nights experiment was performed in a randomized order with an interval of 7 days: one night with normal sleep and another with whole night SD. During the night with normal sleep, participants were asked to sleep a minimum of 6 hours in the Sleep Lab and we assessed their sleep quality using whole-night PSG. Next morning, we acquired high density EEG/fMRI data during rest (5 minutes) and when participants performed three cognitive tasks: MCT (5 minutes) (24), N-back (8 minutes) (23), and ANT (13 minutes)(22). T1-weighted anatomical MRI was also obtained. During the whole night with SD, an investigator stayed overnight with the participant to ensure that the participant did not fall asleep. The next morning, we acquired the same set of imaging data. The participant was then asked to take a nap inside the MRI scanner for a maximum of one hour, during which we continued acquiring high density EEG/fMRI data. After one hour of nap, we woke up the participant and acquired the same set of rest and task imaging data. Finally, a T1-weighted anatomical MRI was obtained.

### Participants

We recruited 53 volunteers according to the following inclusion criteria: participants aged between 18 to 30 years, healthy and good sleepers. Among them, 34 volunteers were selected as eligible after self-report assessment of sleep quality and ruling out exclusion criteria (see *SI Appendix*). Selected participants were then involved in a three nights protocol **Fig. 1A**), in the sleep laboratory of PERFORM Centre at Concordia University. After the first night assessment, 3 participants were excluded due to periodic leg movements and sleep apnea. 5 participants were later excluded because of several unexpected technical issues and 6 participants withdrew before the end of the experiment. Finally, 20 participants completed the whole protocol.

### EEG-fMRI Preprocessing

We used BrainVision Analyzer 2 software (Brain Products, Munich, Germany) for EEG preprocessing and denoising from MR related artefacts. After preprocessing of EEG data acquired during nap inside the MRI scanner, we conducted the scoring of sleep-stages by visual inspection using Wonambi software, a Python-based toolbox for EEG visualization and analysis (https://github.com/wonambi-python/wonambi). A detailed presentation of EEG data preprocessing for this study was reported in (12, 25). Sleep-stage scoring on EEG segments from data simultaneously acquired during the nap was used to identify fMRI runs corresponding to every sleep stage for each subject. Only the EEG segments corresponding to a continuous stable sleep stage of 5 minutes or more were considered for further fMRI analyses. To ensure that we analyzed and compared the same number of frames per state, we trimmed each continuous segment in sleep data, such that only the first 120 time-frames were selected from the runs, if the length of time-course was longer than 120 time- frames (5 min). fMRI data were then preprocessed using the Neuroimaging Analysis Toolkit (NIAK) version 0.13.0, with the MINC Tool Kit (48) (see details in *SI Appendix*). After scrubbing time frames exhibiting excessive motion (frame displacement *>* 0.5*mm*) (27), the remaining motion artefacts were further estimated and removed within SPARK. Slow time drifts were removed by applying a high pass filter using discrete cosine basis functions to ensure a cutoff of 0.01*Hz*. Several runs were excluded when more than 35% of the total time frames were removed by scrubbing. Finally, we selected a set of 5-minute consecutive N2 data, followed by 5-minute N3 data.

### Sparsity-based analysis of k-hubness (SPARK) and HSI estimation

A detailed presentation of SPARK analysis was reported in (18, 19). For each preprocessed fMRI run, we estimated the organization of functional connector hubs in individual brain networks at the voxel level using SPARK. Using SPARK, the whole-brain spatio-temporal activity in fMRI can be decomposed into *N* resting state networks, estimating *N* time-course characteristics (temporal features) and the corresponding *N* spatial maps. The collection of *N* temporal features is called a subject-specific dictionary. SPARK is based on a data-driven sparse general linear model (GLM) (21), which represents the BOLD signal measured in each voxel as a weighted linear combination of *k* temporal features. As the main specificity of SPARK method, *k*_*v*_ is the level of sparsity specific to this voxel, indicating that only a small number of networks, among those estimated *N* networks, are actually involved to describe the fMRI time course of this specific voxel, within a sparse GLM framework (21). Voxel-level k-hubness is typically a discrete number ranging from 1 to 10 networks. Hubs are then defined as brain voxels that are involved in more than one functional network (i.e., *k*_*v*_ *>* 1) that are overlapping in time and space. Region-level *k*-hubness (*k*_*r*_) was also estimated for each region, defined using the Yeo-7 functional networks (29). Then, *HSI*_*v*_ for each voxel was estimated by the ratio of *k*_*r*_ and *k*_*v*_. Finally, ⟨*HSI*⟩_*r*_ for each region *r* was obtained by averaging *HSI*_*v*_ across the voxels belonging to the region *r. ⟨HSI*⟩ _*w*_ was obtained by averaging across the voxels in the whole cortex belonging to the Schaefer-100 atlas(49). Before computing *k*_*v*_ and *k*_*r*_, we performed a visual inspection of *N* network maps to manually classify and exclude the maps that were still contaminated by physiological noise, such as cardiac artifacts, respiratory artifacts or subject in-scanner motion, according to the manual classification criteria proposed in Griffanti et al (50). In average, 15.34*±*3.59 atoms were estimated from SPARK across subjects before manual denoising. 3.97 *±* 1.53 noisy atoms were discarded by the manual denoising procedure.

### Voxel-wise between-state comparisons

To compare our hubness measures (e.g., *k*_*v*_ or *HSI*_*v*_) between different brain states (WR, N2, N3), we performed voxel-wise two sample permutation tests, therefore taking into account inter-subject variability through non-parametric approaches. Individual hubness maps (i.e. maps of *k*_*v*_ over the whole brain) were collected at two different brain states A and B as groups. Student’s *t*-statistic of *k*_*v*_ was computed for each voxel. A null distribution (composed of random groups A and B) was generated by randomly sampling subjects across 10,000 permutations. Finally, an empirical p-value was estimated for each voxel, and adjusted using False Discovery Rate (FDR) correction (see *SI Appendix*).

### Comparison of HSI with FCR

To compare our hub measures with FCR (12, 16), we used a baseline cortical parcellation obtained using the Schaefer-100 atlas(49) and two levels of functional brain networks: Yeo-17 and Yeo 7 networks (29, 49). Using the method described in Cross et al. (12), FCR was computed at the whole cortex level (*FCR*_*w*_) and regional level (*FCR*_*r*_) for seven regions defined using Yeo-7 networks. Since FCR could not be estimated at the voxel level, we proposed an estimation of HSI at the region (*r*) level. When comparing HSI and FCR at the region level, in order to take into account the variance of HSI within one region *r* instead of relying only on one averaged value, we introduced a resampling procedure to obtain 1,000 bootstrap replications of ⟨*HSI*⟩_*r*_ values for each region *r*. A bootstrap replication ⟨*HSI*⟩ _*r*(*boot*)_ of 1000 times was generated by resampling a new set of *n* parcels with replacement within each of the *r* regions, before computing the average. For each replication ⟨*HSI*⟩ _*r*(*boot*)_, the corresponding correlation coefficient (*ρ*_(*boot*)_) and p-value (*p*_(*boot*)_) between ⟨*HSI*⟩ _*r*(*boot*)_ and *FCR*_*r*_ estimated from the same Yeo-7 network *r* were calculated.

## Contributions

KL, YW, TTD, and CG designed research; AJ, ÜA, NC, CA, BF, KL, YW, FR, FP, AAP, AN and MU curated data; KL and YW performed research, analyzed and visualized data; KL, YW and CG wrote the original draft. All authors reviewed the paper.

## ACKNOWLEDGEMENTS

This research was supported by the Natural Sciences and Engineering Research Council of Canada (TDV) and the Canada Foundation for Innovation (TDV). KL was supported by the Wallace H. Coulter Foundation and National Institute on Alcohol Abuse and Alcoholism (NIAAA)/NIH/DHHS grants (5P50AA012870 and 5U01MH121766). The MRI compatible high-density EEG device (Magstim EGI) and data acquisition were made possible through an internal grant from PERFORM center and the Faculty of Arts and Science of Concordia University (CG). TDV is also supported by the Canadian Institutes of Health Research (MOP 142191, PJT 153115, PJT 156125 and PJT 166167), the Fonds de Recherche du Québec – Santé and Concordia University. CG is supported by Natural Sciences and Engineering Research Council of Canada Discovery grants as well as the Canadian Institutes of Health Research (PJT-159948 and MOP-133619) and the Fonds de Recherche du Québec, Nature et Technology (research team grant). UA was supported by postdoctoral fellowships from Savoy Foundation and The Fonds de recherche du Quebec—Sante (FRQS). TDV has received consultant fees from Eisai, Paladin Labs and Jazz Pharmaceuticals. The funders had no role in study design, data collection and analysis, decision to publish, or preparation of the manuscript.

## References

1. Richard B Berry, Rita Brooks, Charlene E Gamaldo, Susan M Harding, Carole Marcus, Bradley V Vaughn, et al. The aasm manual for the scoring of sleep and associated events. Rules, Terminology and Technical Specifications, Darien, Illinois, American Academy of Sleep Medicine, 176(2012):7, 2012.

2. Luigi De Gennaro and Michele Ferrara. Sleep spindles: an overview. Sleep medicine reviews, 7(5):423–440, 2003.

3. Thien Thanh Dang-Vu, Maxime Bonjean, Manuel Schabus, Mélanie Boly, Annabelle Darsaud, Martin Desseilles, Christian Degueldre, Evelyne Balteau, Christophe Phillips, An-dré Luxen et al. Interplay between spontaneous and induced brain activity during human non-rapid eye movement sleep. Proceedings of the National Academy of Sciences, 108 (37):15438–15443, 2011.

4. Thien Thanh Dang-Vu, Manuel Schabus, Martin Desseilles, Geneviève Albouy, Mélanie Boly, Annabelle Darsaud, Steffen Gais, Géraldine Rauchs, Virginie Sterpenich, Gilles Vandewalle, et al. Spontaneous neural activity during human slow wave sleep. Proceedings of the National Academy of Sciences, 105(39):15160–15165, 2008.

5. Chen Song, Melanie Boly, Enzo Tagliazucchi, Helmut Laufs, and Giulio Tononi. fmri spectral signatures of sleep. Proceedings of the National Academy of Sciences, 119(30): e2016732119, 2022.

6. Nina E Fultz, Giorgio Bonmassar, Kawin Setsompop, Robert A Stickgold, Bruce R Rosen, Jonathan R Polimeni, and Laura D Lewis. Coupled electrophysiological, hemodynamic, and cerebrospinal fluid oscillations in human sleep. Science, 366(6465):628–631, 2019.

7. Beverly Setzer, Nina E Fultz, Daniel EP Gomez, Stephanie D Williams, Giorgio Bonmassar, Jonathan R Polimeni, and Laura D Lewis. A temporal sequence of thalamic activity unfolds at transitions in behavioral arousal state. Nature communications, 13(1):5442, 2022.

8. Stephen M Smith, Peter T Fox, Karla L Miller, David C Glahn, P Mickle Fox, Clare E Mackay, Nicola Filippini, Kate E Watkins, Roberto Toro, Angela R Laird, et al. Correspondence of the brain’s functional architecture during activation and rest. Proceedings of the national academy of sciences, 106(31):13040–13045, 2009.

9. Silvina G Horovitz, Allen R Braun, Walter S Carr, Dante Picchioni, Thomas J Balkin, Masaki Fukunaga, and Jeff H Duyn. Decoupling of the brain’s default mode network during deep sleep. Proceedings of the National Academy of Sciences, 106(27):11376–11381, 2009.

10. Jack A De Havas, Sarayu Parimal, Chun Siong Soon, and Michael WL Chee. Sleep deprivation reduces default mode network connectivity and anti-correlation during rest and task performance. Neuroimage, 59(2):1745–1751, 2012.

11. Philipp G Sämann, Renate Wehrle, David Hoehn, Victor I Spoormaker, Henning Peters, Carolin Tully, Florian Holsboer, and Michael Czisch. Development of the brain’s default mode network from wakefulness to slow wave sleep. Cerebral cortex, 21(9):2082–2093, 2011.

12. Nathan E Cross, Florence B Pomares, Alex Nguyen, Aurore A Perrault, Aude Jegou, Makoto Uji, Kangjoo Lee, Fatemeh Razavipour, Obaï Bin Ka’b Ali, Umit Aydin, et al. An altered balance of integrated and segregated brain activity is a marker of cognitive deficits following sleep deprivation. PLoS biology, 19(11):e3001232, 2021.

13. Victor I Spoormaker, Pablo M Gleiser, and Michael Czisch. Frontoparietal connectivity and hierarchical structure of the brain’s functional network during sleep. Frontiers in neurology, 3:80, 2012.

14. Victor I Spoormaker, Manuel S Schröter, Pablo M Gleiser, Katia C Andrade, Martin Dresler, Renate Wehrle, Philipp G Sämann, and Michael Czisch. Development of a large-scale functional brain network during human non-rapid eye movement sleep. Journal of Neuroscience, 30(34):11379–11387, 2010.

15. Marcello Massimini, Fabio Ferrarelli, Reto Huber, Steve K Esser, Harpreet Singh, and Giulio Tononi. Breakdown of cortical effective connectivity during sleep. Science, 309(5744):2228–2232, 2005.

16. Mélanie Boly, Vincent Perlbarg, Guillaume Marrelec, Manuel Schabus, Steven Laureys, Julien Doyon, Mélanie Pélégrini-Issac, Pierre Maquet, and Habib Benali. Hierarchical clustering of brain activity during human nonrapid eye movement sleep. Proceedings of the National Academy of Sciences, 109(15):5856–5861, 2012.

17. Guillaume Marrelec, Pierre Bellec, Alexandre Krainik, Hugues Duffau, Mélanie Pélégrini-Issac, Stéphane Lehéricy, Habib Benali, and Julien Doyon. Regions, systems, and the brain: hierarchical measures of functional integration in fmri. Medical image analysis, 12(4): 484–496, 2008.

18. Kangjoo Lee, Jean-Marc Lina, Jean Gotman, and Christophe Grova. Spark: Sparsity-based analysis of reliable k-hubness and overlapping network structure in brain functional connectivity. NeuroImage, 134:434–449, 2016.

19. Kangjoo Lee, Hui Ming Khoo, Jean-Marc Lina, François Dubeau, Jean Gotman, and Christophe Grova. Disruption, emergence and lateralization of brain network hubs in mesial temporal lobe epilepsy. NeuroImage: Clinical, 20:71–84, 2018.

20. Kangjoo Lee, Corey Horien, David O’Connor, Bronwen Garand-Sheridan, Fuyuze Tokoglu, Dustin Scheinost, Evelyn MR Lake, and R Todd Constable. Arousal impacts distributed hubs modulating the integration of brain functional connectivity. NeuroImage, 258:119364, 2022.

21. Kangjoo Lee, Sungho Tak, and Jong Chul Ye. A data-driven sparse glm for fmri analysis using sparse dictionary learning with mdl criterion. IEEE Transactions on Medical Imaging, 30(5):1076–1089, 2010.

22. Jin Fan, Bruce D McCandliss, John Fossella, Jonathan I Flombaum, and Michael I Posner. The activation of attentional networks. Neuroimage, 26(2):471–479, 2005.

23. Michael WL Chee and Wei Chieh Choo. Functional imaging of working memory after 24 hr of total sleep deprivation. Journal of Neuroscience, 24(19):4560–4567, 2004.

24. Kenneth L Lichstein, Brant W Riedel, and Stephanie L Richman. The mackworth clock test: A computerized version. The Journal of psychology, 134(2):153–161, 2000.

25. Makoto Uji, Nathan Cross, Florence B Pomares, Aurore A Perrault, Aude Jegou, Alex Nguyen, Umit Aydin, Jean-Marc Lina, Thien Thanh Dang-Vu, and Christophe Grova. Data-driven beamforming technique to attenuate ballistocardiogram artefacts in electroencephalography–functional magnetic resonance imaging without detecting cardiac pulses in electrocardiography recordings. Human Brain Mapping, 42(12):3993–4021, 2021.

26. Nathan Cross, Casey Paquola, Florence B Pomares, Aurore A Perrault, Aude Jegou, Alex Nguyen, Umit Aydin, Boris C Bernhardt, Christophe Grova, and Thien Thanh Dang-Vu. Cortical gradients of functional connectivity are robust to state-dependent changes following sleep deprivation. Neuroimage, 226:117547, 2021.

27. Jonathan D Power, Anish Mitra, Timothy O Laumann, Abraham Z Snyder, Bradley L Schlaggar, and Steven E Petersen. Methods to detect, characterize, and remove motion artifact in resting state fmri. Neuroimage, 84:320–341, 2014.

28. Alexander A Borbély, Fritz Baumann, Daniel Brandeis, Inge Strauch, and Dietrich Lehmann. Sleep deprivation: effect on sleep stages and eeg power density in man. Electroencephalography and clinical neurophysiology, 51(5):483–493, 1981.

29. BT Thomas Yeo, Fenna M Krienen, Jorge Sepulcre, Mert R Sabuncu, Danial Lashkari, Marisa Hollinshead, Joshua L Roffman, Jordan W Smoller, Lilla Zöllei, Jonathan R Polimeni, et al. The organization of the human cerebral cortex estimated by intrinsic functional connectivity. Journal of neurophysiology, 2011.

30. Nathalie Tzourio-Mazoyer, Brigitte Landeau, Dimitri Papathanassiou, Fabrice Crivello, Octave Etard, Nicolas Delcroix, Bernard Mazoyer, and Marc Joliot. Automated anatomical labeling of activations in spm using a macroscopic anatomical parcellation of the mni mri single-subject brain. Neuroimage, 15(1):273–289, 2002.

31. Sophie Achard, Chantal Delon-Martin, Petra E Vértes, Félix Renard, Maleka Schenck, Francis Schneider, Christian Heinrich, Stéphane Kremer, and Edward T Bullmore. Hubs of brain functional networks are radically reorganized in comatose patients. Proceedings of the National Academy of Sciences, 109(50):20608–20613, 2012.

32. Thomas T Liu and Maryam Falahpour. Vigilance effects in resting-state fmri. Frontiers in neuroscience, 14:511047, 2020.

33. Tobias Kaufmann, Torbjørn Elvsåshagen, Dag Alnæs, Nathalia Zak, Per Ø Pedersen, Linn B Norbom, Sophia H Quraishi, Enzo Tagliazucchi, Helmut Laufs, Atle Bjørnerud, et al. The brain functional connectome is robustly altered by lack of sleep. Neuroimage, 127:324–332, 2016.

34. Aude Jegou, Manuel Schabus, Olivia Gosseries, Brigitte Dahmen, Geneviève Albouy, Mar-tin Desseilles, Virginie Sterpenich, Christophe Phillips, Pierre Maquet, Christophe Grova, et al. Cortical reactivations during sleep spindles following declarative learning. Neuroimage, 195:104–112, 2019.

35. Emily Cowan, Anli Liu, Simon Henin, Sanjeev Kothare, Orrin Devinsky, and Lila Davachi. Sleep spindles promote the restructuring of memory representations in ventromedial pre-frontal cortex through enhanced hippocampal–cortical functional connectivity. Journal of Neuroscience, 40(9):1909–1919, 2020.

36. Pan Feng, Benjamin Becker, Tingyong Feng, and Yong Zheng. Alter spontaneous activity in amygdala and vmPFC during fear consolidation following 24 h sleep deprivation. NeuroImage, 172:461–469, 2018. ISSN 1053-8119. doi: 10.1016/j.neuroimage.2018.01.057.

37. Yanjing Wang, Cimin Dai, Yongcong Shao, Chuan Wang, and Qianxiang Zhou. Changes in ventromedial prefrontal cortex functional connectivity are correlated with increased risk-taking after total sleep deprivation. Behavioural brain research, 418:113674, 2022.

38. James M Shine, Patrick G Bissett, Peter T Bell, Oluwasanmi Koyejo, Joshua H Balsters, Krzysztof J Gorgolewski, Craig A Moodie, and Russell A Poldrack. The dynamics of functional brain networks: integrated network states during cognitive task performance. Neuron, 92(2):544–554, 2016.

39. Peter T Fox and Karl J Friston. Distributed processing; distributed functions? Neuroimage, 61(2):407–426, 2012.

40. Matthew A Wilson and Bruce L McNaughton. Reactivation of hippocampal ensemble memories during sleep. Science, 265(5172):676–679, 1994.

41. Reto Huber, M Felice Ghilardi, Marcello Massimini, and Giulio Tononi. Local sleep and learning. Nature, 430(6995):78–81, 2004.

42. Bryce A Mander, Shawn M Marks, Jacob W Vogel, Vikram Rao, Brandon Lu, Jared M Saletin, Sonia Ancoli-Israel, William J Jagust, and Matthew P Walker. β-amyloid disrupts human nrem slow waves and related hippocampus-dependent memory consolidation. Nature neuroscience, 18(7):1051–1057, 2015.

43. Bruce Luber, Jason Steffener, Adrienne Tucker, Christian Habeck, Angel V Peterchev, Zhi-De Deng, Robert C Basner, Yaakov Stern, and Sarah H Lisanby. Extended remediation of sleep deprived-induced working memory deficits using fmri-guided transcranial magnetic stimulation. Sleep, 36(6):857–871, 2013.

44. Jeffrey S Durmer and David F Dinges. Neurocognitive consequences of sleep deprivation. In Seminars in neurology, volume 25, pages 117–129. Copyright© 2005 by Thieme Medical Publishers, Inc., 333 Seventh Avenue, New …, 2005.

45. Hans PA Van Dongen, Amy M Bender, and David F Dinges. Systematic individual differences in sleep homeostatic and circadian rhythm contributions to neurobehavioral impairment during sleep deprivation. Accident Analysis & Prevention, 45:11–16, 2012.

46. Ed Bullmore and Olaf Sporns. Complex brain networks: graph theoretical analysis of structural and functional systems. Nature reviews neuroscience, 10(3):186–198, 2009.

47. Kangjoo Lee, Jie Lisa Ji, Markus Helmer, John D Murray, John H Krystal, and Alan Anticevic. A framework for advancing mechanistic neuro-behavioral biomarkers in psychiatry. Biological Psychiatry, 99(10):896–908, 2025.

48. Pierre Bellec, Felix Carbonnell, Vincent Perlbarg, Claude Lepage, Oliver Lyttelton, Valdmir Fonov, Andrew Janke, Jussi Tohka, and Alan Evans. A neuroimaging analyses kit for matlab and octave. In Human Brain Mapping HBM 2011 17th Annual Meeting of the Organization on Human Brain Mapping, Quebec City, Canada, June 26–30, 2011, pages 1–5. Organization on Human Brain Mapping, 2011.

49. Alexander Schaefer, Ru Kong, Evan M Gordon, Timothy O Laumann, Xi-Nian Zuo, Avram J Holmes, Simon B Eickhoff, and BT Thomas Yeo. Local-global parcellation of the human cerebral cortex from intrinsic functional connectivity mri. Cerebral cortex, 28(9):3095–3114, 2018.

50. Ludovica Griffanti, Gwenaëlle Douaud, Janine Bijsterbosch, Stefania Evangelisti, Fidel Alfaro-Almagro, Matthew F Glasser, Eugene P Duff, Sean Fitzgibbon, Robert Westphal, Davide Carone, et al. Hand classification of fmri ica noise components. Neuroimage, 154: 188–205, 2017.

